# Tailored haemodynamic response function increases detection power of fMRI in awake dogs (*Canis familiaris*)

**DOI:** 10.1101/2020.03.12.987537

**Authors:** Magdalena Boch, Sabrina Karl, Ronald Sladky, Ludwig Huber, Claus Lamm, Isabella C. Wagner

## Abstract

Functional magnetic resonance imaging (fMRI) of awake and unrestrained dogs (*Canis familiaris*) has been established as a novel opportunity for comparative neuroimaging, promising important insights into the evolutionary roots of human brain function and cognition. However, data processing and analysis pipelines are often derivatives of methodological standards developed for human neuroimaging, which may be problematic due to profound neurophysiological and anatomical differences between humans and dogs. Here, we explore whether dog fMRI studies would benefit from a tailored dog haemodynamic response function (HRF). In two independent experiments, dogs were presented with different visual stimuli. BOLD signal changes in the visual cortex during these experiments were used for (a) the identification and estimation of a tailored dog HRF, and (b) the independent validation of the resulting dog HRF estimate. Time course analyses revealed that the BOLD signal in the primary visual cortex peaks significantly earlier in dogs compared to humans, while being comparable in shape. Deriving a tailored dog HRF significantly improved the model fit in both experiments, compared to the canonical HRF used in human fMRI. Using the dog HRF yielded significantly increased activation during visual stimulation, extending from the occipital lobe, to the caudal parietal cortex, the bilateral temporal cortex, and into bilateral hippocampal and thalamic regions. In sum, our findings provide robust evidence for an earlier onset of the dog HRF in a visual stimulation paradigm, and suggest that using such an HRF will be important to increase fMRI detection power in canine neuroimaging. By providing the parameters of the tailored dog HRF and related code, we encourage and enable other researchers to validate whether our findings generalize to other sensory modalities and experimental paradigms.

**Highlights:**

- Dog fMRI typically uses human HRF, but underlying neurophysiology might differ
- V1 BOLD signal peaked earlier in dogs than predicted by the human HRF
- Tailored dog HRF improved model fit when tested with independent data
- Whole-brain comparisons confirmed increased detection power for tailored dog HRF
- Dog fMRI will benefit from increased detection power of tailored HRF

## 1 Introduction

Animal research involving domesticated dogs (*Canis familiaris*) yield important insights into non-invasive comparative neuroscience (Andics, Gácsi, Faragó, Kis, & Miklósi, 2014; Bunford, Andics, Kis, Miklósi, & Gácsi, 2017; Fitch, Huber, & Bugnyar, 2010), and allows researchers to study the neural correlates of cognitive abilities, i.e., how dogs perceive or process their environment (e.g. Andics & Miklósi, 2018; Thompkins, Deshpande, Waggoner, & Katz, 2016 for review). For example, recent work has used functional magnetic resonance imaging (fMRI) to study the neural representations during auditory stimulation or lexical processing (Andics et al., 2016, 2014; Prichard et al., 2019; Prichard, Cook, Spivak, Chhibber, & Berns, 2018), face perception (Cuaya, Hernández-Pérez, & Concha, 2016; Dilks et al., 2015; Hernández-Pérez, Concha, & Cuaya, 2018; Szabó et al., 2020), olfactory processing (Berns, Brooks, & Spivak, 2015; Jia et al., 2014), sense for numeracy (Aulet et al., 2019), jealousy (Cook, Prichard, Spivak, & Berns, 2018) and reward processing (Berns, Brooks, & Spivak, 2012; Berns, Brooks, Spivak, & Levy, 2017; Berns, Brooks, & Spivak, 2013; Cook, Prichard, Spivak, & Berns, 2016; Cook, Spivak, & Berns, 2014; Prichard, Chhibber, Athanassiades, Spivak, & Berns, 2018) in dogs. So far, dog fMRI studies have relied on methodological standards originally developed for human (f)MRI, but it has been proposed that hardware as well as data analysis approaches tailored to dogs might be more suitable (Huber & Lamm, 2017). Although the majority of fMRI pre-processing steps are readily transferable from humans to dogs (i.e., slice timing correction, realignment, smoothing), humans and dogs might differ in many aspects other than apparent differences in neuroanatomy (Hecht et al., 2019; Horschler et al., 2019; Schoenebeck & Ostrander, 2013), such as differences in vascular and neuronal physiology. Here, we critically examined the state of the art in canine neuroimaging methodology and aimed at optimizing data processing and analysis pipelines to improve fMRI sensitivity and specificity.

fMRI-based neuroimaging commonly uses a general linear model (GLM) to describe voxel-wise haemodynamic response time courses by convolving the regressors of the experimental conditions with a haemodynamic response function (referred to as “human HRF” throughout the text). This typically involves a double-gamma function to account for the delayed peak at approx. 5 s after stimulus onset and the post-stimulus undershoot (Friston, Fletcher, et al., 1998; Friston et al., 1995; Friston, Jezzard, & Turner, 1994; Worsley & Friston, 1995). So far, canine neuroimaging studies have used the standard human HRF (e.g., Andics et al., 2014; Cuaya, Hernández-Pérez, & Concha, 2016), a model of the human HRF based on a single gamma function (e.g., Cook et al., 2014; Dilks et al., 2015), or a Fourier basis set (Aguirre et al., 2007). However, assumptions about the (canonical) human HRF shape and its temporal dynamics might not apply in dogs. An accurate HRF model is crucial, as even minor deviations can lead to substantial loss of power (Handwerker, Ollinger, & D’Esposito, 2004), thus not only reducing the chance of detecting true effects but also increasing the likelihood for false-positive results (Lindquist, Meng Loh, Atlas, & Wager, 2009) and inflated effect sizes (e.g., Ioannidis, 2005; Simmons, Nelson, & Simonsohn, 2011). Additionally, fMRI studies with small sample sizes are often considered underpowered (Button et al., 2013; Cremers, Wager, & Yarkoni, 2017; Poldrack et al., 2017; Simmons et al., 2011), which is a ubiquitous problem in canine research due to the complexity of the experiments (median of approx. 12.5 dogs, although it is increasing). Under these circumstances, it is particularly crucial to test whether the BOLD response in dogs is adequately captured with the canonical human HRF, or some variations of it.

The shape of the human HRF has been discussed extensively since its adoption in fMRI data analysis (Aguirre, Zarahn, & D’Esposito, 1998; Boynton, Engel, Glover, & Heeger, 1996; Glover, 1999). Numerous factors causing HRF variability have been identified, e.g., developmental changes (Arichi et al., 2012), and clinical conditions (Ford, Johnson, Whitfield, Faustman, & Mathalon, 2005). A frequent approach to account for potential HRF variability within a participant sample (used twice in a dog sample, Jia et al., 2014, 2016) is to add temporal and/or dispersion derivatives (TDD) along with the HRF regressor when applying the GLM, used to calculate a so-called informed basis set (Friston, Fletcher, et al., 1998; Friston, Josephs, Rees, & Turner, 1998; Henson, Price, Rugg, Turner, & Friston, 2002). Despite the increased flexibility in the model, the basis function depends on prior knowledge about the average shape of the underlying BOLD signal, which is currently not available in canine neuroscience research.

Previous studies using invasive recordings indeed demonstrated that the HRF varies across mammalian species. In comparison to humans, the HRF was shown to peak earlier in rats (De Zwart et al., 2005; Lambers et al., 2020; Silva, Koretsky, & Duyn, 2007) and mice (Chen et al., 2020), while the HRF in macaque monkeys appears similar (Baumann et al., 2010; Goense & Logothetis, 2008; Koyama et al., 2004; Logothetis, Pauls, Augath, Trinath, & Oeltermann, 2001; Nakahara, Hayashi, Konishi, & Miyashita, 2002; Patel, Cohen, Baker, Snyder, & Corbetta, 2018). Deviations from the human HRF in terms of shape and temporal dynamics seem to decrease in species with closer common ancestry to humans (Upham, Esselstyn, & Jetz, 2019) and with increasing absolute brain size (e.g., Roth & Dicke, 2005 for review). Considering the variations across species and potential differences in underlying neurophysiology, it seems plausible that the human HRF might deviate from the average BOLD signal in dogs. However, precise conclusions are currently not possible, as systematic investigations of the BOLD signal have not yet been performed in dogs.

Here, we aimed to close this gap and used non-invasive fMRI in awake dogs that were specifically trained for this approach. In two independent experiments, we used different visual stimulation experiments and a step-wise analysis approach to establish and validate our results, respectively. In the first experiment, dogs viewed a flickering checkerboard interspersed with a baseline condition (flickering checkerboard experiment, experiment 1). The experiment employed a block design, aimed at achieving a robust measure of the average BOLD signal in the primary visual cortex (V1). Based on the resulting V1 BOLD signal data, we identified and estimated a tailored dog HRF, compared its model fit to the one based on using the human HRF, and differences in whole-brain activation between the two HRFs. We also tested if adding time and dispersion derivatives to the human HRF could sufficiently account for potential deviations of the dog from the human HRF. Data from a second experiment, which had employed an event-related visual stimulation design (face processing experiment, experiment 2), were then used to validate the results from the flickering checkerboard experiment. We opted for visual stimulation as the V1 can be easily located (see e.g., Langley & Grünbaum, 1890; Marquis, 1934; Uemura, 2015; Wing & Smith, 1942), thus ameliorating the problem of a common three-dimensional coordinate system in canines. Finally, to encourage reproducibility, we openly share our data and provide a detailed description of the processing and analysis pipeline (see also for similar challenges on reproducibility in human fMRI: Carp, 2012b, 2012a; Nichols et al., 2017; Poldrack et al., 2017, 2008). Together, our results provide a first investigation on whether the human HRF model appropriately fits the average BOLD signal in dogs and whether estimating a novel dog HRF can increase fMRI specificity and detection power.

## 2 Materials and methods

### 2.1 Sample

Seventeen pet dogs participated in the flickering checkerboard experiment (experiment 1; 10 females, age range = 3-11 years, mean = 7.24 years, *SD* = 2.33 years); consisting of 12 border collies, 2 Australian shepherds, 1 border collie Australian shepherd mix, 1 Labrador retriever and 1 mixed-breed dog (weight range = 15-27 kg, mean = 19.67 kg, *SD* = 3.87). A subsample of fourteen dogs also participated in the face processing experiment (experiment 2; 8 females, age range = 3-11 years, mean = 7.21 years, *SD* = 2.46 years) in the same or max. two months apart; consisting of 10 border collies, 1 Labrador retriever, 1 Australian shepherd, 1 border collie Australian shepherd mix and 1 mixed-breed dog (weight range = 15-27 kg, mean = 19.25 kg, *SD* = 4.03).

All dogs passed an initial medical examination concerning eyesight and general health. The human caregivers gave written informed consent to their dogs’ participation and did not receive any monetary compensation. The dogs were fully awake and unrestrained, and were able to exit the MR scanner at any time. To achieve this, they received extensive training prior to the MRI sessions in order to habituate them to the MRI environment (see Karl, Boch, Virányi, Lamm, & Huber, 2019 for a detailed description of the training procedure, and Berns & Cook, 2016; Strassberg, Waggoner, Deshpande, & Katz, 2019 for similar procedures). The study was approved by the institutional ethics and animal welfare commission in accordance with Good Scientific Practice (GSP) guidelines and national legislation at the University of Veterinary Medicine Vienna (ETK-06/06/2017), based on a pilot study conducted at the University of Vienna. The current study complies with the ARRIVE Guidelines (Kilkenny, Browne, Cuthill, Emerson, & Altman, 2010).

### 2.2 Experimental setup

#### Preparation

Together with the dog trainer, the dog entered the MR scanner room wearing earplugs and an additional head bandage to secure optimal earplug positioning and to enhance noise protection. The dog then accessed the scanner bed via a custom-made ramp and positioned the head inside the coil, seated in sphinx position (Figure 1A). The dog trainer then moved the dog into the scanner bore and visual tasks were presented using an MR-compatible computer screen placed at the end of the scanner bore (32 inch). Additionally, we used the camera of an eye-tracker (Eyelink 1000 Plus, SR Research, Ontario, Canada) to ensure that the dogs stayed awake, did not close their eyes during stimulus onsets, and to monitor overall movement (*N* = 5 dogs in experiment 2 were not monitored due to later implementation of the camera). The dog trainer remained in the MR-scanner room throughout the entire scan session but left the dog’s visual field before task onset. The majority of the dogs first participated in the flickering checkerboard experiment followed by the face processing experiment in a subsequent MR-session (Figure 1B). Data acquisition was aborted if the dog moved extensively (as observed using eye-tracking, see above) or if the dog exited the scanner bore during the task. Data collection was then repeated within the same or a subsequent session, depending on the dog’s motivation. Following the scan session, we evaluated the realignment parameters and re-invited the dog to repeat the experiment in a subsequent session if head motion exceeded a threshold of 3 mm (Figure 1C). On average, two scan sessions were necessary to complete the experiment below the motion threshold for both experiments; 12 out of 17 dogs and 9 out of 14 dogs succeeded in their first scan session for experiment 1 and experiment 2, respectively. After completing a run, the dog exited the MR scanner and received a food reward.

**Figure 1.**
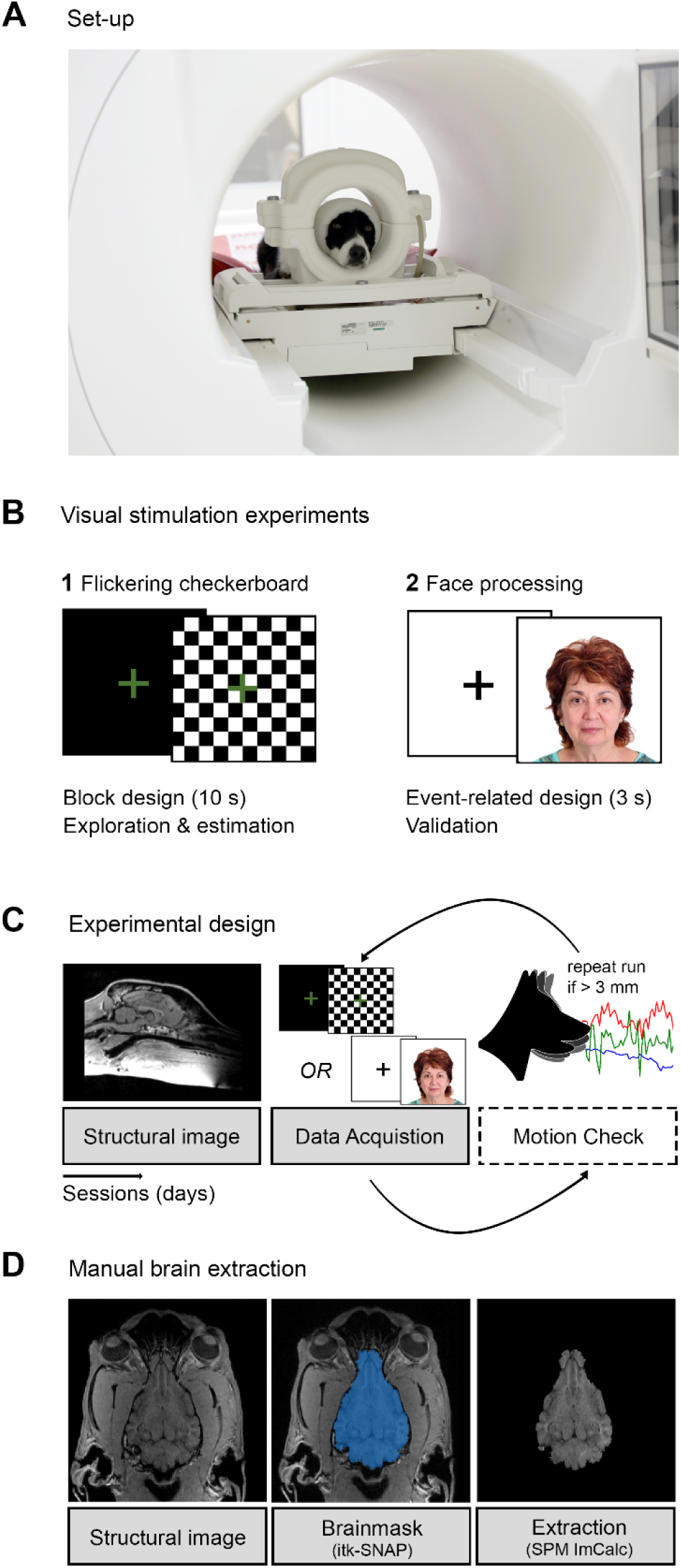
Overview of experimental approach to explore the average BOLD signal in dogs and estimate a tailored dog haemodynamic response function (HRF). (**A**) All dogs were trained to position their head in a 15-channel human knee coil and to stay motionless during data acquisition. (**B**) We acquired data in two different visual stimulation experiments. In (1), we extracted the average primary visual cortex (V1) BOLD signal using data from a flickering checkerboard experiment, and estimated a tailored dog haemodynamic response function (HRF). We compared this dog HRF to the canonical human HRF, and the human HRF with time and dispersion derivatives (TDD). Second, in (2), we validated the results using a face processing experiment, whose data served as an independent test data set. (**C**) Structural scans were acquired in a session prior to functional data acquisition of the visual stimulation experiments; functional tasks were acquired in separate sessions. Movement parameters were assessed after successful completion of a task. If motion exceeded 3 mm, we repeated the task in additional sessions. (**D**) We created individual tailor-made brain masks using itk-SNAP (Yushkevich et al., 2006) to skull-strip the structural images and consequently improve co-registration and normalization of the canine neuroimaging data.

#### Flickering checkerboard experiment (experiment 1)

The task used in this experiment alternated between blocks of visual stimulation (flickering checkerboard covering the whole screen and green cross in the centre for 10 s) and a visual baseline with a green cross presented on a black screen for 10 s. The total task duration was 2.2 min, including six blocks of visual stimulation and 6 blocks of baseline in a fixed order, starting with the visual baseline condition (see Figure 1B). We chose this experiment for the dog HRF estimation based on the fact that a block design can be expected to be more robust and predictable, even if the human and dog HRFs and the actual BOLD signal time courses differ.

#### Face processing experiment (experiment 2)

The task for experiment 2 alternated between short events of visual stimulation (3 s clips of varying conditions, showing smooth transitions between two facial expressions from different human models, all on white background; 500 × 500 pixels) and a visual baseline with a black cross on a white screen jittered between 5-7 s (see Figure 1B). Within the scope of the present methodological study, we focused on visual responses compared to baseline, irrespective of the different conditions (results of this will be reported elsewhere). The total task comprised 60 trials of visual stimulation split in two runs. Each run took 5 min with a short break outside the MR scanner if both runs were acquired in the same session.

### 2.3 MRI data acquisition

Data were collected using a 3T Siemens Skyra MR-system using a 15-channel coil developed for structural imaging of the human knee. Functional imaging data for both tasks were obtained from 24 axial slices (interleaved acquisition; descending order, covering the whole brain) using a 2-fold multiband-accelerated echo planar imaging (EPI) sequence and a voxel size of 1.5 × 1.5 × 2 mm^3^ (TR/TE = 1000/38 ms, field of view (FoV) = 144 x 144 x 58 mm^3^, flip angle = 61°, 20% gap). The task from experiment 1 (flickering checkerboard experiment) consisted of a single run comprising 134 scans, and the task employed in experiment 2 (face processing experiment) comprised two runs of 270 scans each. The dogs had multiple attempts to complete the task in case of excessive head motion (see 2.2. experimental design). The structural image was obtained using a voxel size of 0.7 mm isotropic (TR/TE = 2100/3.13 ms, FoV = 230 × 230 × 165 mm^3^) and was acquired in a prior scan session, separated from the functional imaging sessions.

### 2.4 Data processing and statistical analysis

#### 2.4.1 MRI data preprocessing

All imaging data was analysed using SPM12 (https://www.fil.ion.ucl.ac.uk/spm/software/spm12/) and Matlab 2014b (MathWorks) (see Figure 2 for an overview of the workflow). After slice timing correction (referenced to the middle slice, Sladky et al., 2011) and image realignment, the functional images were manually reoriented to match the orientation of the canine breed-averaged template (Nitzsche et al., 2017) with the rostral commissure as a visual reference using the SPM module “*Reorient images / Set origin”*. We then manually skull-stripped the structural image using an individual binary brain mask for each dog, created using itk-SNAP (Yushkevich et al., 2006). Based on preliminary analyses, skull-stripping canine imaging data proved to be essential for successful automatic co-registration. This way, the co-registration algorithm successfully detects brain borders, not incorrectly relying on large muscles that surround the dog brain but have similar image intensity (see Figure 1D). The structural image, the individual binary brain mask, and the functional imaging data were then co-registered to the mean image of each run. Next, the structural image was segmented (“*Old Segmentation*” module of SPM12) into grey matter, white matter, and cerebrospinal fluid, using the tissue probability maps provided by the canine breed-averaged template (Nitzsche et al., 2019). We then normalized (using the “*Old Normalization*” module of SPM12) the functional and structural imaging data, along with the individual binary brain mask. Lastly, functional images were resliced (1.5 mm isotropic) and smoothed using a 3 mm Gaussian kernel (full-width-at-half-maximum, FWHM).

**Figure 2.**
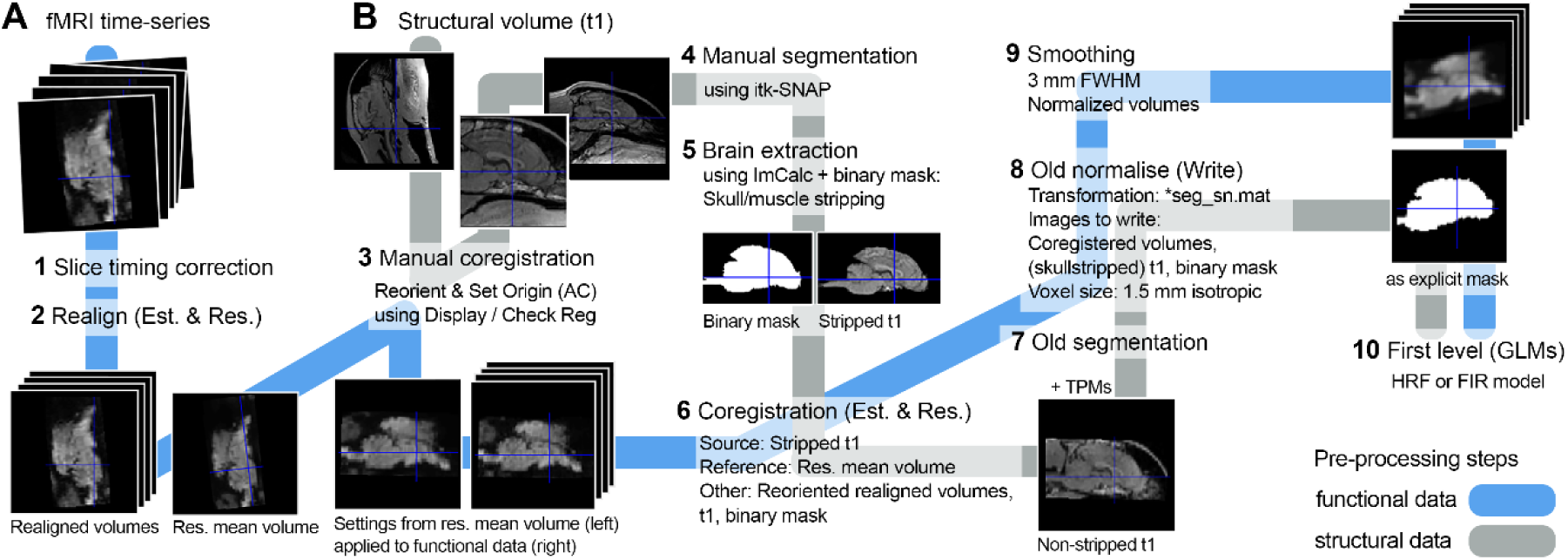
Schematic description of the tailored data processing workflow for the canine neuroimaging data including (**A**) functional images and (**B**) the structural image. Exemplary structural and functional images as well as binary mask are from one dog in the sample; tissue probability maps (TPMs) are from the canine breed-averaged atlas (Nitzsche et al., 2019). Numbers, in bold, describe the sequence of processing steps. Est., estimate; Res., resliced, GLM, general linear model.

To additionally account for head motion, we performed motion scrubbing by calculating the scan-to-scan motion for each dog, referring to the framewise displacement (FD) between the current scan *t* and its preceeding scan *t*-1. For each scan that exceeded the FD threshold of 0.5 mm, we entered an additional motion regressor to the first-level GLM design matrix (Power, Barnes, Snyder, Schlaggar, & Petersen, 2012; Power et al., 2014). For the checkerboard experiment (experiment 1), on average 7.8% of the scans were removed (∼10/134 scans, ranging from 0 to 36 scans). For the face processing experiment (experiment 2), on average 3.5% (run 1) and 5.5% (run 2) scans were removed (run 1: ∼ 10/270 scans; run 2: ∼ 15/270 scans; ranging from 0 to 52 across runs).

#### 2.4.2 Template normalization

We attempted to provide a unified coordinate system by combining two available templates, (1) based on a canine breed-average (Nitzsche et al., 2019) combined with (2) the normalized labels from another canine template based on a single male Golden Retriever (Czeibert, Andics, Petneházy, & Kubinyi, 2019). First, we segmented (“Old Segmentation”) the structural template (Czeibert et al., 2019) using the tissue probability maps provided by the breed-averaged template (Nitzsche et al., 2019). Then, we normalized (“Old Normalization”) both the structural template and the NIfTI-file containing the atlas labels.

### 2.5 fMRI data analysis

We now provide an overview of the analysis approach followed by more details on each analysis step in the following section (see also Figure 3). For the exploratory investigation of the average BOLD signal and estimation of the tailored dog HRF, we first analysed activation changes in V1 during experiment 1 (contrast flickering checkerboard > visual baseline) in the following steps: (1) we extracted the average V1 time course of the BOLD signal employing a finite impulse response (FIR) model (exploration and estimation analysis step 1, extraction V1 BOLD signal); (2) we estimated a tailored dog HRF based on the FIR data above (exploration and estimation analysis step 2, dog HRF estimation); (3) we then compared the human HRF with the dog HRF using model fit analysis and Wilcoxon signed ranks tests (exploration and estimation analysis step 3, model fit comparison). Then, to expand comparisons to the whole-brain, (4) we performed first-level analysis using the human HRF, the human HRF with time and dispersion derivatives and the tailored dog HRF (exploration and estimation analysis step 4, first-level GLMs) and (5) analysed neuroimaging data on a group-level along with paired *t*-tests (exploration and estimation analysis steps 5, group-level activation comparisons)

**Figure 3.**
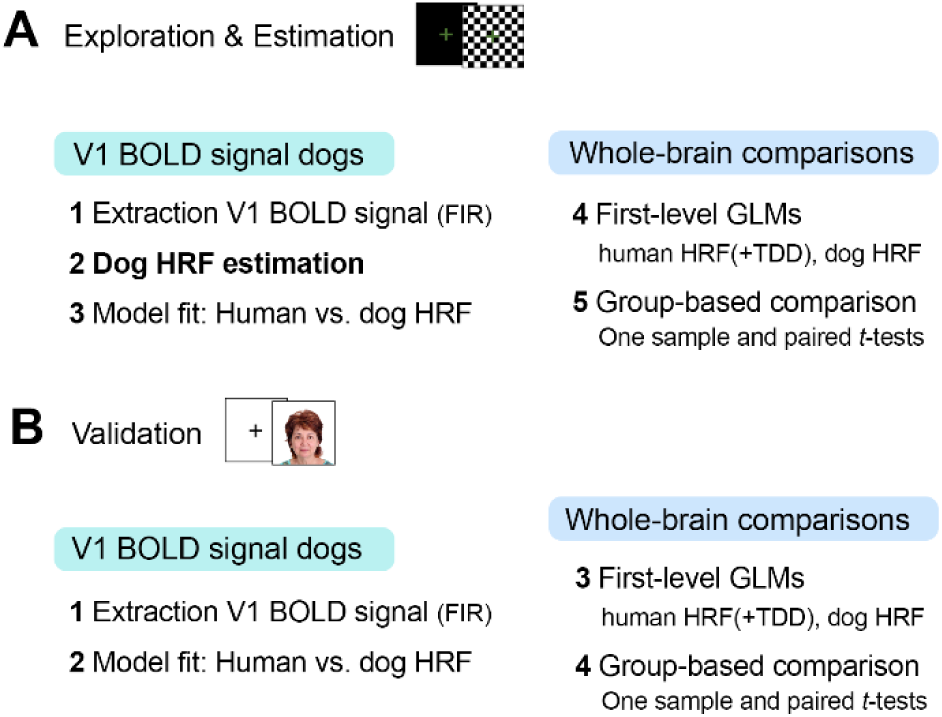
Overview on analyses underpinning exploration of the average V1 BOLD signal in dogs, estimation of the tailored dog haemodynamic response function (HRF), and validation of that HRF in a second independent data set. (**A**) Data from the flickering checkerboard experiment served for the exploratory and estimation analysis to (1) to extract the average V1 BOLD signal in dogs and visually compare it to the human HRF model using a finite impulse response (FIR) model, (2) estimate a tailored dog HRF based on the empirical data and (3) compare model fits of the human and dog HRF in the visual cortex. On the whole-brain level, (4) we then performed first-level analyses using the human HRF, the human HRF along with time and dispersion derivatives (TDD) and the tailored dog HRF to (5) perform whole-brain group comparisons using one sample and paired *t*-tests across HRF models. (**B**) Results from (A) where then validated using the data from the face processing experiment as an independent validation data set. All analysis steps were as above, except for dog HRF estimation. GLM, general linear model; BOLD, Blood Oxygenation Level Dependent

Next, to validate the results from experiment 1, which revealed an earlier peak of the V1 BOLD signal in dogs, we cross-validated them by analysing V1 activation changes during the face processing experiment (contrast faces > visual baseline), using a similar but modified approach: (1) we extracted the average time course of the V1 BOLD signal during the face processing experiment using a FIR model (validation step 1, extraction V1 BOLD signal); (2) we compared the HRF models based on their model fit and using Wilcoxon signed ranks tests (validation step 2, model comparison); (3) we performed univariate activation analysis using the human HRF, the human HRF along with time and dispersion derivatives (TDD), and the dog HRF (validation step 3, first-level GLMs); lastly, (4) we performed group activation analyses along with paired *t*-tests (validation step 4, group-level activation comparisons).

#### 2.5.1 Exploration and estimation analysis: Flickering checkerboard experiment (experiment 1)

##### Step 1: Extraction average V1 BOLD signal

We used a finite impulse response (FIR) model to measure the average V1 time course of the BOLD signal in dogs. This flexible approach makes minimal assumptions about the shape of the BOLD signal and thus results in independent response estimates for a predetermined number of time bins (in the present case, one time bin per TR). We estimated FIRs covering the visual stimulation blocks (starting at stimulus onset (0 s) until 10 s after stimulus offset), yielding a duration of 20 s that was divided in 20 time bins (TR = 1 s). We then extract the average V1 time course, based on V1 coordinates obtained from the group-based comparison using the human HRF (exploratory and estimation analysis step 5; see also Table 1, section “human HRF”), using (a) a 4 mm sphere placed around the local maximum of the cluster that covered the occipital lobe (Figure 4A) and (b) expanding over V1 as determined by our atlas labels (Czeibert et al., 2019; Nitzsche et al., 2019). Finally, we extracted each dog’s average BOLD time series and calculated the time course of activation induced by the visual stimulation block across all dogs.

**Table 1.** Flickering checkerboard experiment: Task-related activation during visual stimulation.

| Contrast, brain region & HRF | Coordinates<br>(breed-averaged template) |  |  | z-value | cluster size |
| --- | --- | --- | --- | --- | --- |
|  | x | y | z |  |  |
| **Human HRF:** Flickering checkerboard > visual baseline (k = 14) |  |  |  |  |  |
| L caudal splenial gyrus (O) | -1 | -29 | 16 | 5.71 | 610 |
| L hippocampus (T) | -9 | -18 | 1 | 4.47 | 15 |
| **Human HRF+TDD:** Flickering checkerboard > visual baseline (k = 10) |  |  |  |  |  |
| L caudal splenial gyrus (O) | 1 | -26 | 19 | 6.05 | 246 |
| R medial suprasylvian gyrus (T) | 13 | -20 | 18 | 4.61 | 11 |
| **Dog HRF:** Flickering checkerboard > visual baseline (k = 15) |  |  |  |  |  |
| R caudal splenial gyrus (O) | 1 | -27 | 18 | 6.23 | 823 |
| L medial suprasylvian gyrus (T) | -16 | -18 | 19 | 4.82 | 30 |
| L caudal suprasylvian gyrus (T) | -19 | -24 | 7 | 4.17 | 18 |
| L hippocampus (T) | -9 | -17 | 3 | 4.08 | 23 |
| R hippocampus (T) | 8 | -15 | 6 | 4.04 | 19 |
| **Human HRF+TDD > human HRF:** Flickering checkerboard > visual baseline (k = 10) |  |  |  |  |  |
| L caudal splenial gyrus (O) | -3 | -30 | 16 | 5.58 | 175 |
| **Human HRF > dog HRF:** Flickering checkerboard > visual baseline (k = 14) |  |  |  |  |  |
| L insular cortex (T) | -18 | -11 | -2 | 4.58 | 14 |
| **Dog HRF > human HRF: Flickering checkerboard > visual baseline ($k = 14$)** |  |  |  |  |  |
| R caudal splenial gyrus (O) | 1 | -27 | 18 | 5.78 | 316 |
| R medial suprasylvian gyrus (T) | 17 | -18 | 21 | 4.71 | 54 |
| L medial suprasylvian gyrus (T) | -16 | -20 | 21 | 4.17 | 14 |
| **Human HRF+TDD > dog HRF: Flickering checkerboard > visual baseline ($k = 10$)** |  |  |  |  |  |
| L caudal splenial gyrus (O) | -3 | -30 | 16 | 6.00 | 162 |

Effects were tested for significance with a cluster-defining threshold of $p < 0.001$ and a cluster probability of $p < 0.05$ FWE-corrected for multiple comparisons. Critical cluster sizes ( $k$ ) are reported along with the results for each one sample $t$ -test, each haemodynamic response (HRF) model, and for the paired-sample $t$ -tests between all HRF models. The first local maximum within each cluster is reported; coordinates represent the location of peak voxels and refer to the canine breed-averaged template (Nitzsche et al., 2019), the template along with another single dog template (Czeibert et al., 2019) served to determine anatomical nomenclature for all tables. TDD, time and dispersion derivatives; O, occipital lobe; T, temporal lobe; L, left; R, right.

**Figure 4.**
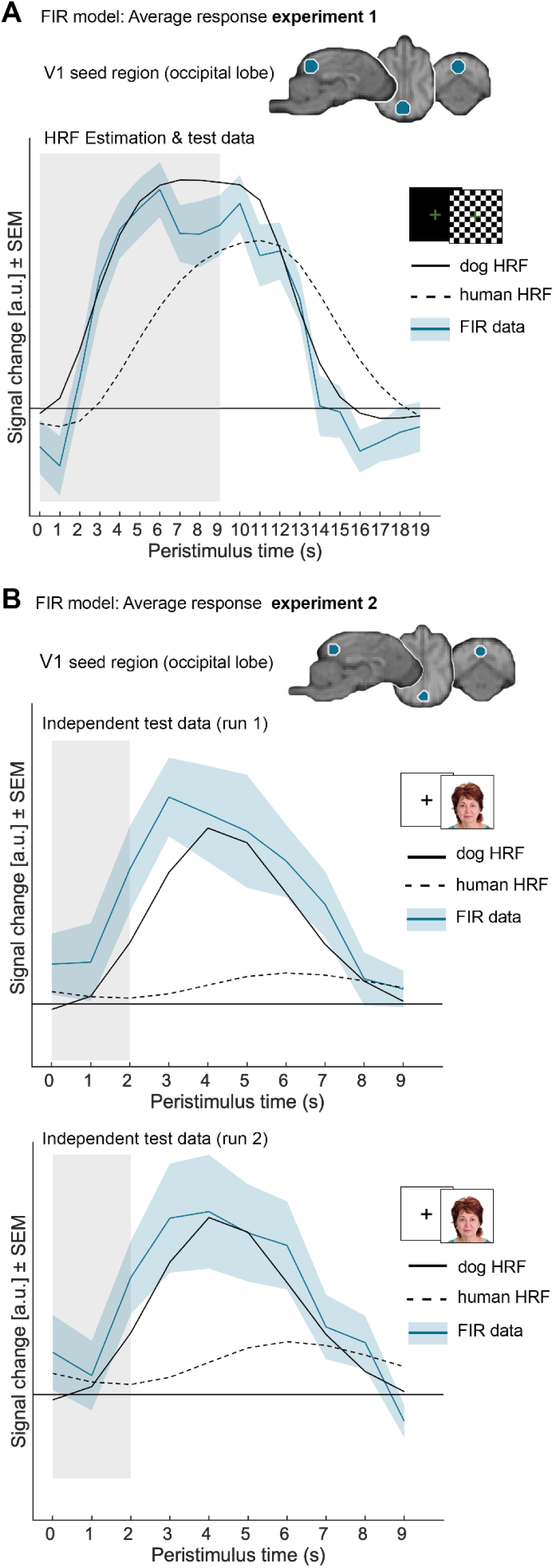
Visual comparison reveals an earlier peak of the BOLD signal in dogs than modelled by the canonical human haemodynamic response function (HRF) for both independent data sets leading to estimation of tailored dog HRF. After calculating the finite impulse response (FIR) models, we extracted individual response estimates from the maximal response in primary visual cortex (V1) using (**A**) the standard human HRF for the flickering checkerboard experiment (exploration and estimation analysis, step 5; x = −1, y = −29, z = 16, 4 mm) and (**B**) the standard human HRF along with time and dispersion derivatives for the face processing experiment (validation analysis, step 4; x = −1, y = −29, z = 19, 4 mm). Based on the extracted data, we calculated the response to the visual stimulation across trials averaged for all dogs for both (**A**) the flickering checkerboard experiment and (**B**) the face processing experiment (both runs separately). The dog HRF was estimated based on the FIR results from the flickering checkerboard experiment (exploration and estimation analysis, step 2), the face processing experiment served as an independent test data set to validate the results derived from the exploration and estimation analysis. For illustration purposes, the dog and human HRF’s were scaled by the parameter estimates from the respective GLM’s.

##### Step 2: Estimation of the dog HRF

Based on the results from step 1, which upon visual inspection revealed the need for a tailored dog HRF with earlier onset, we estimated a new parametrization for SPM’s canonical HRF, yielding a tailored dog HRF model. The *spm_hrf* function uses seven optional parameters to specify the shape of the HRF: the delay of the response (relative to onset, *p*_*1*_ = 6 s), the delay of the undershoot (relative to onset, *p*_*2*_ = 16 s), the dispersion of the response (*p*_*3*_ = 1), the dispersion of undershoot (*p*_*4*_ = 1), the ratio of the response to the undershoot (*p*_*5*_ = 6), the onset (*p*_*6*_ = 0 s), and the length of the kernel (*p*_*7*_ = 32 s). We used MATLAB’s *fminsearch* function, a multidimensional unconstrained nonlinear minimization method, to optimize the model fit of the regression analysis (*R*^*2*^-statistics of MATLAB’s *regress* function) by varying the values of *p*_*1*_, *p*_*2*_, *p*_*5*_, *p*_*6*_. The assumed plausible ranges for the haemodynamic parameters were: *p*_*1*_ = [1 10 s], *p*_*2*_ = [1 20 s], *p*_*5*_ = [1 10 s], *p*_*6*_ = [0 5 s], and the regression analysis was identical to a standard SPM first-level analysis (see above, step 1). We chose not to deviate from the well-established default-values for response (*p*_*3*_) or undershoot dispersion (*p*_*4*_), or the overall kernel length (*p*_*7*_) to prevent overfitting.

##### Step 3: Model fit comparison

We then calculated the individual single-subject *R*^*2*^-statistics of each GLM with the different HRF parameters and compared the model fit to the extracted V1 BOLD signal between human and dog HRF using a Wilcoxon signed ranks test.

##### Step 4: Human HRF

Using the GLM approach implemented in SPM12, we estimated contrast images for each dog that reflected task-related activation (contrast checkerboard > baseline). The first-level design matrix of each dog contained a task regressor modelling visual stimulation, time-locked to the onset of each block (duration 10 s) and convolved with the human (canonical) HRF. The six realignment parameters along with regressors modelling framewise displacement (see above) were added to the design matrix to account for head motion. Normalized, and individually created binary masks (see above and Figure 2) were used as explicit masks and a high-pass filter with a cut-off at 128 s was applied.

##### Human HRF+TDD

Next, to account for variability (Friston, Fletcher, et al., 1998; Friston, Josephs, et al., 1998; Henson et al., 2002), we added temporal and dispersion derivatives (TDD) to the human HRF. The visual stimulation regressor was thus convolved with the human HRF along with its TDD. This resulted in three regression parameter estimates consisting of: (1) the human canonical HRF 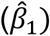, (2) the time derivative 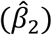, and (3) the dispersion derivative 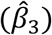. We then combined all three regressors to form one “derivative boost (H)”-regressor per dog (Calhoun, Stevens, Pearlson, & Kiehl, 2004; Lindquist et al., 2009): 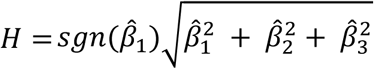.

##### Dog HRF

Next, we set up a first-level model (same settings as previously) including the data that was now estimated and convolved using the estimated dog HRF (step 3, human HRF).

##### Step 5: Group-level activation comparison

To test for activation differences during visual stimulation on a group-level, we implemented one sample *t*-tests for each HRF model (steps 1, 2, 5; contrasting flickering checkerboard > baseline; H-regressor for TDD model), as well as paired sample *t*-tests (checkerboard > baseline). Unless stated otherwise, significance was determined using cluster-level inference with a cluster-defining threshold *p* < 0.001 and a cluster probability of *p* < 0.05 family-wise error (FWE) corrected for multiple comparisons. Cluster extent was calculated using the SPM extension “CorrClusTh.m” (by Thomas Nichols, University of Warwick, United Kingdom, and Marko Wilke, University of Tübingen, Germany; http://www2.warwick.ac.uk/fac/sci/statistics/staff/academicresearch/nichols/scripts/spm/).

#### 2.5.2 Validation analysis: Face processing experiment (experiment 2)

Independent data obtained during the face processing experiment (experiment 2) were then used to validate the exploratory results and to compare all three HRF models.

##### Step 1: Extraction average V1 BOLD signal

Similar to above (exploration and estimation analysis, step 1) we used a finite impulse response (FIR) model to extract the individual BOLD signal time courses, but defined 10 time bins starting at the stimulus onset (0 s) until 7 s after stimulus offset (each time bin had a duration of 1 s = length of TR). We then placed a 4 mm sphere around the local maxima of the cluster encompassing the V1, and used the coordinates emerging from the human HRF+TDD model (validation analysis, step 5; Table 2 section “*human HRF+TDD*”) since the human HRF did not survive the significance threshold (Figure 4B).

**Table 2.** Face processing experiment: Task-related activation during visual stimulation.

| Contrast, brain region & HRF | Coordinates<br>(breed-averaged template) |  |  | z-value | cluster size |
| --- | --- | --- | --- | --- | --- |
|  | x | y | z |  |  |
| **Human HRF+TDD:** Visual stimulation > visual baseline ($k = 21$) | | | | | |
| R lateral olfactorial gyrus (T) | 13 | 3 | -4 | 4.65 | 21 |
| R caudal marginal gyrus (O) | 1 | -29 | 19 | 4.29 | 26 |
| **Dog HRF:** Visual stimulation > visual baseline ($k = 14$) | | | | | |
| R medial suprasylvian gyrus (T) | 16 | -23 | 19 | 4.88 | 255 |
| R rostral ectosylvian gyrus (T) | 17 | -8 | 10 | 4.54 | 53 |
| R caudal splenial gyrus (O) | 2 | -32 | 18 | 4.53 | 130 |
| L caudal composite gyrus (T) | -22 | -20 | -4 | 4.49 | 204 |
| **Dog HRF > human HRF:** Visual stimulation > visual baseline ($k^* = 10$) | | | | | |
| R medial suprasylvian gyrus (T) | 17 | -23 | 18 | 4.19 | 28 |
| L medial suprasylvian gyrus (T) | -18 | -24 | 15 | 3.70 | 24 |
| **Dog HRF > human HRF+TDD:** Visual stimulation > visual baseline ($k^* = 10$) | | | | | |
| R ectomarginal gyrus (P) | 8 | -18 | 19 | 4.04 | 22 |

Effects were tested for significance with a cluster-defining threshold of $p < 0.001$ and a cluster probability of $p < 0.05$ FWE-corrected for multiple comparisons. Critical cluster sizes ( $k$ ) are reported along with the results for each one sample $t$ -test per haemodynamic response function (HRF) model. For the one sample $t$ -test based on the human HRF GLM, no cluster survived the threshold. None of the paired sample $t$ -tests across HRF models survived the critical cluster threshold ( $k$ ), therefore the significance level was lowered to $p < 0.005$ with an arbitrary cluster threshold ( $k^*$ ) of 10 voxels. One paired sample $t$ -test (human HRF vs. human HRF and time and dispersion derivatives (TDD)) did not survive the lowered threshold as well as the contrasts human > dog HRF, human HRF + TDD > dog HRF. The first local maximum within each cluster is reported; coordinates represent the location of peak voxels and refer to the canine breed-averaged template (Nitzsche et al., 2019), the template along with another single dog template (Czeibert et al., 2019) served to determine anatomical nomenclature for all tables. O, occipital lobe; T, temporal lobe; L, left; R, right.

##### Step 2: Model fit comparison

This step was almost identical to above (exploration and estimation analysis, step 3) but was performed based on the FIR data from experiment 2 (validation analysis, step 1).

##### Step 3: Human HRF

Analysis was identical to above (exploration and estimation analysis, step 4 human HRF), but visual stimulation was modelled with one task regressor time locked to the event onset (duration of 3 s), contrasted against visual baseline (contrast faces > baseline).

##### Human HRF+TDD

Analysis was identical to above (exploration and estimation analysis, step 4 human HRF+TDD) using the task regressor from experiment 2 (validation analysis, step 3 human HRF) but resulted in two informed basis sets as this task contained two separate runs. We first calculated the mean of each parameter estimate across both runs (i.e. 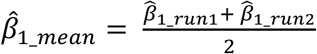) and then, as above, combined all three averaged regressors to one “derivative boost (H)”-regressor per dog.

##### Dog HRF

We defined the same first-level model as described above (validation analysis, step 3 human HRF) but the task regressor was convolved with the newly estimated dog HRF.

##### Step 4: Group-level activation comparison

This step was performed based on the first-level results from experiment 2 but otherwise identical to above (exploration and estimation analysis, step 5).

### 2.6 Data and code availability statement

Unthresholded statistical maps from the exploratory and estimation analysis, the Matlab-based code to estimate the dog HRF and a *spm_my_defaults.m*-script containing the dog HRF parameters have been added as supplementary material and will be available along with the peer-reviewed version of the article.

## 3 Results

### 3.1 Exploration and estimation analysis: Flickering checkerboard experiment (experiment 1)

#### FIR model and dog HRF estimation

To investigate the time course of the BOLD response in dogs, we used a model-free analysis (FIR model, exploration and estimation analysis, step 1). Results suggested a temporal difference between the standard (canonical) human HRF and the average response in our canine sample. Visual inspection of the results revealed an earlier peak after visual stimulation onset compared to convolution using a human HRF and, consequently, an earlier decline and return to baseline (Figure 4A). Therefore, the estimation based on the FIR data (exploration and estimation analysis, step 2) resulted in the following parameter changes to the (canonical) human HRF: a shorter response delay (*p*_*1*_ *=* 4.3 s), a delay of the undershoot (*p*_*2*_ *=* 6.6 s), as well as a lower ratio of the response to the undershoot (*p*_*5*_ *=* 3). This newly estimated dog HRF peaked around 2-3 seconds earlier as compared to the human HRF (Figure 4A).

#### Determining the HRF model fits

*R*^*2*^-statistics of both GLMs calculated individually (main analysis, step 6) revealed a better model fit of the average time course of activation when using the dog HRF, with a mean *R*^*2*^ of 0.64 (*SD* = 0.21), increasing the fit almost two times in comparison to the model using the human HRF (mean *R*^*2*^ = 0.35 (*SD* = 0.20). This substantial increase in explained variance was statistically significant (*z* = 142, *p* = 0.002).

#### Visual activation: Human HRF/human HRF+TDD

Expanding to whole-brain comparisons (exploration and estimation analysis, step 5), we performed standard whole-brain GLM analyses similar to other canine neuroimaging papers (e.g., Andics et al., 2016; Cuaya et al., 2016) and localized visual processing areas by convolving fMRI data with the human HRF (exploration and estimation analysis, step 3 human HRF). Results revealed increased activation within the occipital lobe (V1) and within the left hippocampal area (Table 1, section “human HRF”). When accounting for HRF variability (exploration and estimation analysis, step 3, human HRF+TDD), we found similar activation within V1 during visual stimulation (but only about half the size compared to the human HRF) as well as within the right dorsal temporal lobe (Table 1, section “human HRF+TDD). Additionally, V1 clusters stemming from both analysis types expanded from the occipital lobe to portions of the parietal and right temporal lobe (Figure 5A). Thus, analyses based on the standard human HRF with and without accounting for its variability yielded comparable activation increases in V1 during visual stimulation.

**Figure 5.**
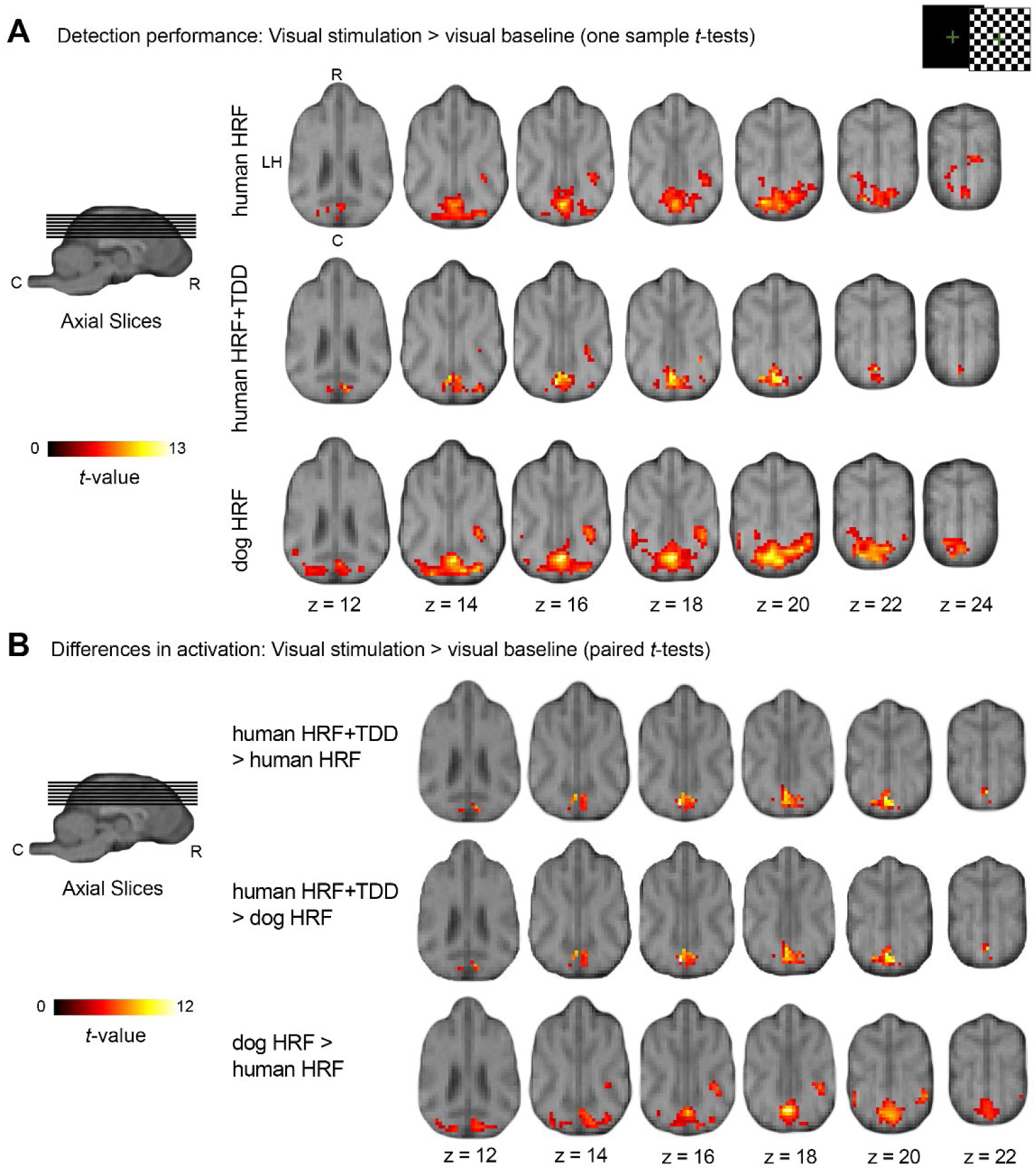
Flickering checkerboard experiment: Comparison of brain activation across haemodynamic response functions (HRF) illustrates increased detection performance using a tailored dog HRF in both primary and higher order visual processing areas (exploratory and estimation analysis). Results are displayed at *p* < 0.05, FWE-corrected at cluster-level, and using a cluster-defining threshold of *p* < .001 (see Table 1) on the mean structural image. Coordinates refer to the canine breed-averaged atlas (Nitzsche et al., 2019). The first axial plane (**A**, first row, left) shows the anatomical locations caudal (C), rostral (R), and left hemisphere (LH); all axial planes displayed have the same orientation. The sagittal plane displays the cut coordinates and the anatomical locations dorsal (D), ventral (V). (**A**) Group-based activation for visual stimulation > baseline (one sample *t*-tests) indicate confirm that the analysis using the dog HRF shows the highest sensitivity for the canine neuroimaging data, with the analysis using the human HRF resulting in smaller and the one using the human HRF combined with time and dispersion derivatives even smaller activation clusters (**B**) Comparisons of visual stimulation > visual baseline contrasts between all three HRF models (paired *t*-tests) resulted in similar significant activation changes in the occipital lobe for the human HRF and time and dispersion (TDD) model in contrast to both the human and dog HRF). Comparing the human HRF and dog HRF revealed stronger activation in the primary visual cortex and temporal regions for the dog HRF compared to the dog HRF and activation in the insular cortex for the reverse contrast (not depicted, see Table 1 for details).

#### Visual activation: Dog HRF

We now report in more detail the brain areas revealing significant activation on a group-level using the tailored dog HRF, since it significantly improved the model fit in the V1 compared to the human HRF (exploration and estimation analysis, steps 2-3). We observed five clusters with stronger activation during visual stimulation compared to baseline (Table 1, section “*dog HRF*”, Figure 5), which is more than double the amount of significant clusters, as well as cluster sizes, compared to the remaining models (main analysis, steps 1, 2; Table 1). The largest cluster expanded from the V1 to bilateral parietal and temporal lobe regions, followed by smaller clusters in the right temporal lobe (see Table 1 and Figure 6 for details).

**Figure 6.**
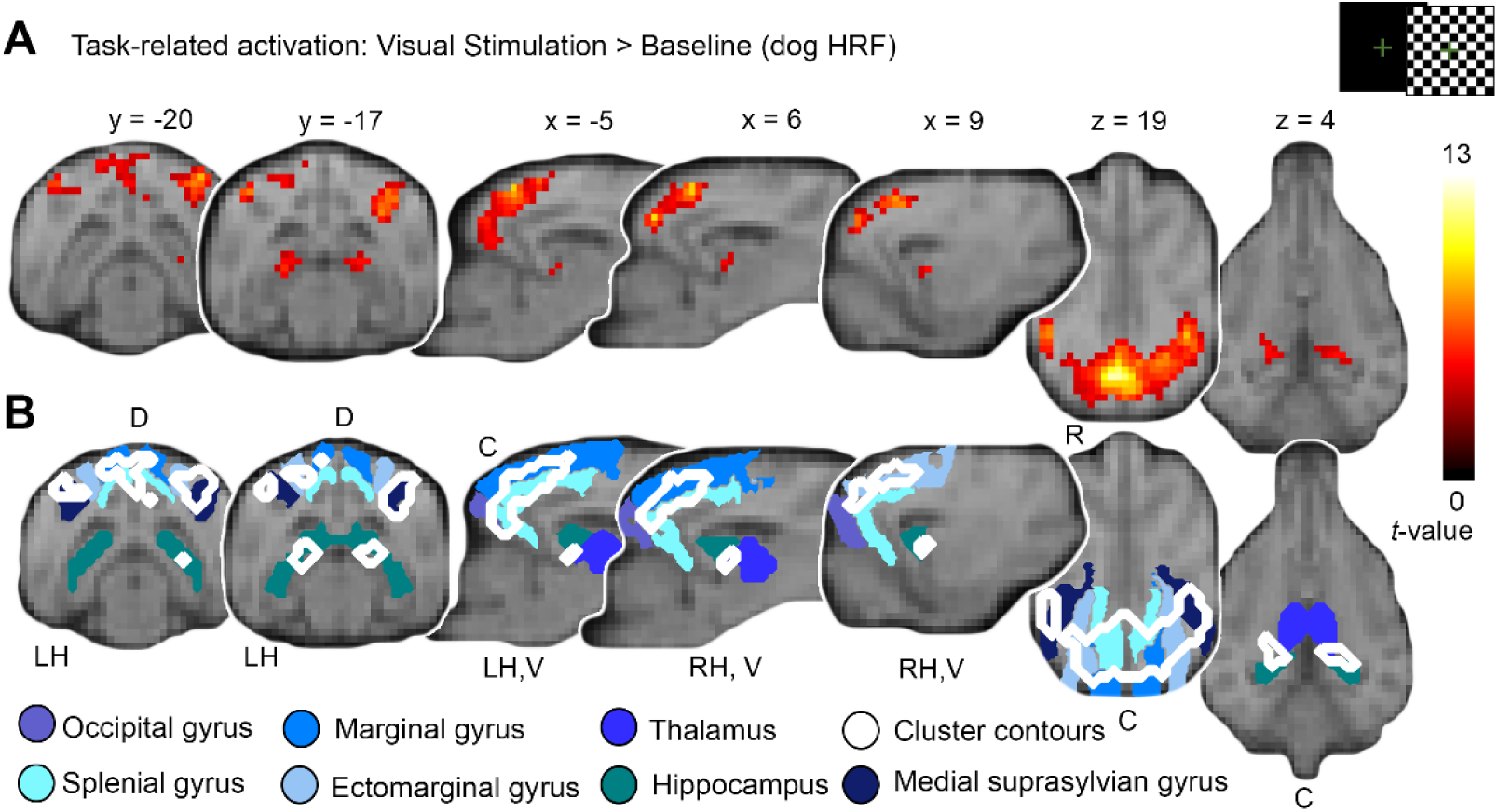
Increasing the detection power by using the tailored dog haemodynamic response function (HRF) in the flickering checkerboard experiment allows detailed description of primary and higher-order visual processing areas. (**A**) Visual stimulation against baseline elicited activation in a large region of the occipital lobe peaking at the rostral occipital lobe expanding to the caudal parietal lobe and bilateral dorsal portions of the temporal lobe. In addition, activation in bilateral hippocampal areas increased in response to visual stimulation compared to baseline. Results are displayed at *p* < 0.05, FWE-corrected at cluster-level, and using a cluster-defining threshold of *p* < .001 (see Table 1, section “dog HRF”), plotted onto the mean structural image. Atlas maps, coordinates and the anatomical nomenclature refer to the canine breed-averaged atlas (Nitzsche et al., 2019) and additional normalized labels from a single-dog based template (Czeibert et al., 2019). Images are accompanied with anatomical locations caudal (C), rostral (R), dorsal (D), ventral (V), left hemisphere (LH) and right hemisphere (RH). (**B**) For easier interpretation of the anatomical structures activated, blue-shaded outlines of anatomical regions are displayed together with contours of activated clusters shown in Panel A.

#### Activation differences during visual stimulation across HRF models

In order to test for whole-brain differences in activation, we compared the human HRF, human HRF+TDD and dog HRF GLMs using paired sample *t*-tests (contrast checkerboard > visual baseline; exploration and estimation analysis, step 5). Results revealed significant clusters for all models. However, the analysis using the dog HRF was the only one that resulted in significant differences in activation both in the V1 and bilateral temporal regions (dog HRF > human HRF); the human HRF+TDD increased activation only in a caudal V1 region (human HRF+TDD > human HRF; human HRF+TDD > dog HRF). In sum, the human HRF revealed to be the least sensitive model (see Figure 5B, Table 1 for details).

### 3.2 Validation: Face processing experiment (experiment 2)

Next, we validated our novel results in an independent data set and compared all three HRF models.

#### FIR model and comparison of HRF model fits

Visual inspection of the average activation time course based on the FIR model (validation analysis, step 4) confirmed the results of the exploratory and estimation analysis, as it again revealed an earlier BOLD signal peak (see Figure 4B). In line with the exploratory results, comparing the average HRF model fits for both runs separately (validation analysis, step 5) revealed that the dog HRF improved the fit by eight times for the first run (human HRF: mean 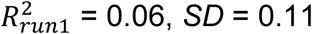; dog HRF: mean 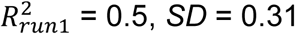) and by almost three times for the second run (human HRF: mean 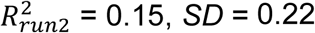; dog HRF: mean 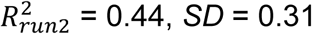). Again, the Wilcoxon signed ranks tests indicated that the dog HRF model fit was significantly higher than the human HRF in both runs (Run 1: *z* = 100, *p* = 0.001; Run 2: *z* = 67, *p* = 0.012), confirming the advantage of using the tailored dog HRF in a data set independent of the dog HRF estimation.

#### Visual activation during visual stimulation across HRF models

In line with the results from the exploratory and estimation analysis, modelling the dog HRF resulted in the highest number of activated clusters with cluster sizes increasing twelve times in comparison to the model including the human HRF+TDD. Furthermore, the dog HRF was the only model that detected activation beyond the V1 in bilateral temporal regions, while none of these withstood the cluster threshold correction when modelling the human HRF (see Table 2, Figure 7A for details; validation analysis, step 5). Performing paired sample *t*-tests between dog HRF, human HRF and human HRF+TDD (validation analysis, step 5) resulted in no significant differences with the initial strict threshold, but lowering the threshold to *p* = 0.005 uncorrected indicated that using the dog HRF improved the sensitivity to detect visual processing areas (see Table 2, Figure 7B for details), thus confirming the exploratory results.

**Figure 7.**
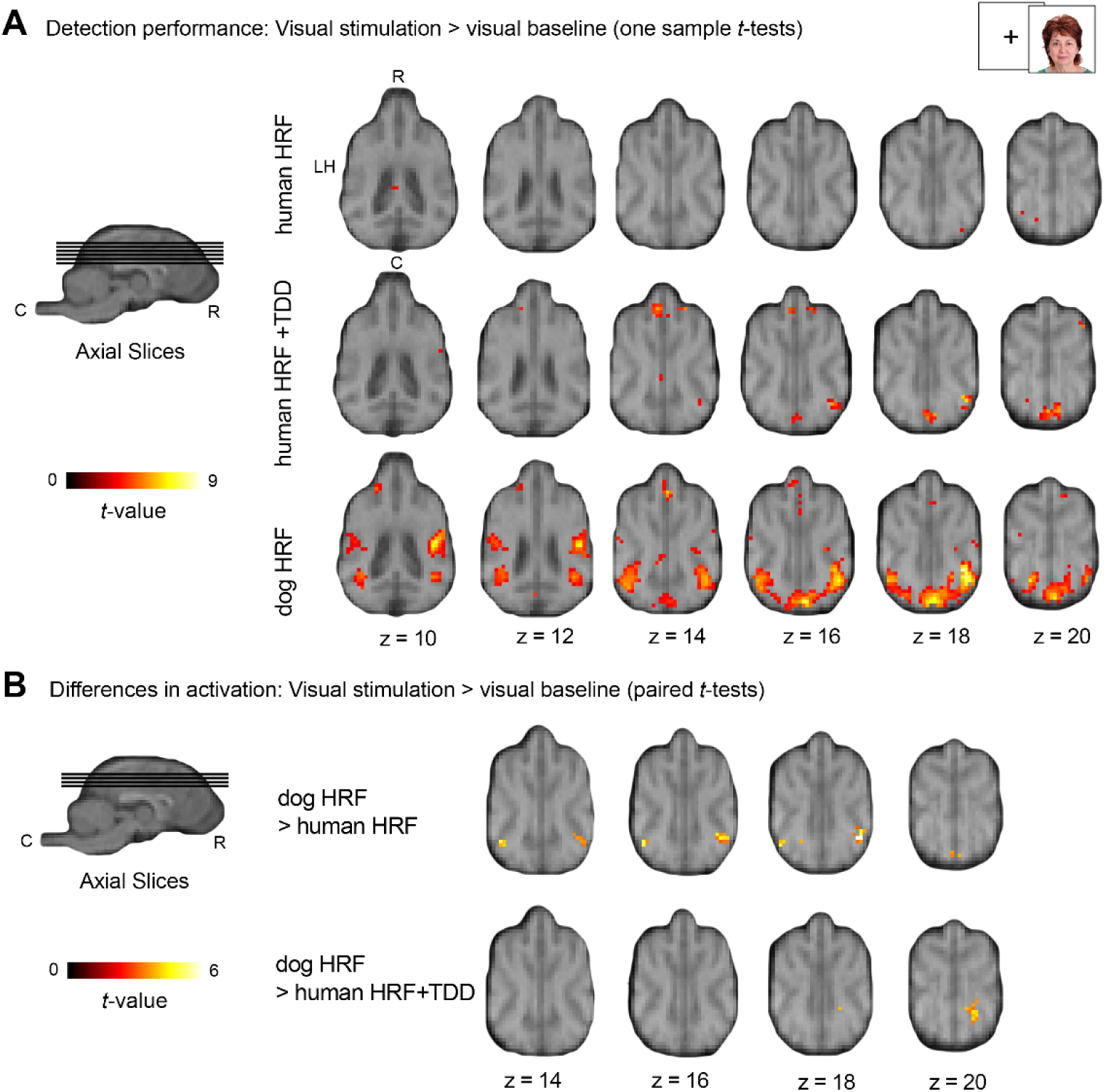
Face processing experiment: Comparison of brain activation in an independent data set confirms increased detection performance using a tailored dog haemodynamic response function (HRF) compared to other HRF models (validation analysis). For display purposes results are displayed at *p* < .005 (for results at *p* < 0.05, FWE-corrected at cluster-level, and a cluster-defining threshold of *p* < .001 see Table 2) on the mean structural image. Coordinates refer to the canine breed-averaged atlas (Nitzsche et al., 2019). The first axial plane (**A**, first row, left) shows the anatomical locations caudal (C), rostral (R) and left hemisphere (LH); all axial planes displayed have the same orientation. The sagittal plane displays the cut coordinates and the anatomical locations dorsal (D), ventral (V). (**A**) Group-based activation for visual stimulation > baseline (one sample *t*-tests) indicate that the human HRF results in almost no activation, the human HRF combined with time and dispersion derivatives (TDD) results in bigger activation clusters and again that the dog HRF shows the highest sensitivity for the canine neuroimaging data. (**B**) Group comparisons of visual stimulation > visual baseline contrasts between all three HRF models. Group-based activation (paired *t*-tests) resulted in trends of activation changes in temporal regions for the dog HRF in comparison to both the human HRF and human HRF + TDD model (see Table 2 for detailed results).

## 4 Discussion

The aim of this study was to explore whether the typically used human haemodynamic response function (HRF) fits the average BOLD signal in dogs and whether detection power for canine neuroimaging data can be improved using a tailored dog HRF. Our results indicate that the human HRF does not fit the average BOLD signal in dogs. We provide initial evidence that the average time course of the V1 BOLD signal in dogs peaks 2-3 s earlier than the human HRF and that the model fit for the primary visual cortex (V1) can be significantly improved using a tailored dog HRF. Expanding to whole-brain activation, the dog HRF again resulted in increased detection power for the dog HRF.

We used two independent visual experiments serving as exploration and estimation analysis (flickering checkerboard experiment, experiment 1) and independent validation sets (face processing experiment, experiment 2). We estimated a tailored dog HRF based on the empirical data from experiment 1, since V1 BOLD signal indicated an earlier peak compared to the human HRF. Following this, we were able to confirm the earlier peak when investigating the V1 BOLD signal in the independent experiment 2. Further, the model fit for the V1 significantly improved (and almost doubled) in experiment 1 and improved between eight (run 1) and almost three (run 2) times in experiment 2 when comparing to the human HRF. Expanding to whole-brain comparisons, our results provide evidence that the human HRF, compared to the tailored dog HRF, resulted in significantly less activation being detected. Fourth, adding time and dispersion derivatives (TDD) led to significantly increased activation in both experiments, but only within occipital areas. Overall, however, the human HRF+TDD was less sensitive in detecting secondary visual areas which resulted in fewer significant clusters during both experiments. These are important findings when considering the small sample sizes in most canine neuroimaging studies. In contrast to human studies, it is more difficult to increase power by increasing the sample size, primarily due to limited availability of canine participants and extensive dog training prior to MR-scanning. Thus, increasing the model fit of the HRF to the average BOLD signal time course is an important alternative tool to further increase the power and therefore increase the reproducibility of future studies.

Our findings are consistent with research in rodents, which suggested that using the human HRF degrades the model fit and, thus, the overall detection performance (Lambers et al., 2020). As in our sample, Lambers and colleagues (2020) observed an earlier peak of the average BOLD signal in rats, proposing differences in brain and vessel size and smaller distances within the brain as potential reasons for the observed patterns. Absolute brain sizes cannot sufficiently explain why the human HRF fits the average BOLD signal in dogs. Although dog brains have a smaller *absolute* size than human and, on average, macaque brains (e.g., DeFelipe, 2011; Yáñez et al., 2005), the dog breeds in our sample seem to have a similar size as rhesus macaques (Horschler et al., 2019). However, *relative* size (brain size/body weight) could potentially explain our findings, since the dog brains in our sample (just as rodent brains) seem to have a smaller relative brain size than humans and rhesus macaques (e.g. Baumann et al., 2010; Logothetis et al., 2001 for average body weight in macaques; Roth & Dicke, 2005 for review). Although evolutionary relationship also seems to correlate with the human HRF across species underlying neurovascular mechanisms remain somewhat unclear. Additionally, skull shapes and sizes also vary *within* dog species (i.e., across different breeds), resulting in substantial variance in underlying neuroanatomy in dogs (Hecht et al., 2019; Horschler et al., 2019; Schoenebeck & Ostrander, 2013). Since our sample was rather homogenous (70% border collies; all mesocephalic skull shapes), we did not have enough variance to test for potential differences between breeds, skull shapes or sizes. Therefore, the average BOLD signal might deviate from the tailored dog HRF across breeds. This could be accounted for by adding time and dispersion derivatives to the dog HRF in future studies.

Our results do not confirm earlier reports (Berns et al., 2012) of a similar time course of the average BOLD signal to the one in humans. Unlike Berns et al. (2012), who reported a comparable time course of activation in dogs, our results suggest that the human HRF does not fit the average time course of the BOLD signal in dogs optimally. However, Berns et al. (2012) studied the subcortical caudate nucleus, while we focused on the cortical BOLD signal in dogs, extracting data from the V1. Previous research in other species, i.e. humans showed that the average BOLD signal time course differs between cortical and subcortical regions (Handwerker et al., 2004; Lewis, Setsompop, Rosen, & Polimeni, 2018). Thus, our findings do not necessarily contradict the results from Berns et al. (2012) but might be related to the different areas analysed, as well as their neural and vascular characteristics.

Overall, our findings provide first evidence that the human HRF in the visual cortex does not optimally fit the HRF observed in dogs. Despite being based on two independent experiments allowing for cross-validation, this evidence should be treated as preliminary, awaiting independent validation in other samples, experimental paradigms, and brain regions. We hope that our approach will encourage future research to test the reproducibility and generalizability of our findings, and to explore whether this could help to increase model fit and detection power in their own canine fMRI datasets. For this reason, we adopted the established and recommended (Carp, 2012b; Nichols et al., 2017; Poldrack et al., 2008) standards from human neuroimaging analyses, provided a detailed description of our workflow and parameters, and made our imaging data and code openly available. Using a simple but salient sensory stimulation experiment also allowed quality assessment of our developed processing pipeline and helped us validate future changes in our pipeline, preventing potentially biased decisions. Additionally, a short (visual) localizer experiment can be used for dog training and getting dogs accustomed to the experimental setup.

Transparent reporting also allows us to build on previous research and facilitates the comparison of results. Based on previous research (e.g., Aguirre et al., 2007; Langley & Grünbaum, 1890; Marquis, 1934; Uemura, 2015; Willis, Quinn, McDonell, Gati, Parent, et al., 2001; Willis, Quinn, McDonell, Gati, Partlow, et al., 2001; Wing & Smith, 1942) research, we are certain about the location of the V1, but less is known about other higher-order visual association areas. Similar to the human and rhesus macaque visual system (e.g., Orban, Van Essen, & Vanduffel, 2004; Tootell, Tsao, & Vanduffel, 2003 for comparative reviews), we found activation within the dorsal visual stream, extending from the occipital lobe to the caudal parietal lobe and the ventral stream, and including bilateral regions in the temporal lobes, bilateral hippocampus and caudal thalamus. We did not find significant activation in the lateral geniculate body (LGB); (1) regarding the small size of the region, detection might require smaller voxel sizes or (2) differences in individual anatomy might have led to anatomical imprecision, atlases based on larger sample size (Nitzsche et al. 2019: based on *N* = 16 dogs) could help disentangle this question. Unfortunately, there is still no agreement on a shared template space; publicly available templates (Czeibert et al., 2019; Datta et al., 2012; Liu et al., 2020; Nitzsche et al., 2019) are not in the same space and vary in orientation and origin, thus coordinates from one template cannot be applied to the other. Taken together, these findings can be a next step to further investigate the visual system for dogs, hopefully aiding future investigations of the visual system in dogs or studies focusing on visual paradigms (e.g., face processing Cuaya et al., 2016; Dilks et al., 2015; Hernández-Pérez et al., 2018; Szabó et al., 2020; Thompkins et al., 2018).

### 4.1 Conclusions

We present first evidence that the average visual-cortical BOLD signal in dogs peaks earlier than the human HRF model. Consequently, the significantly lower model fit suggests that the analysis of canine neuroimaging data using the human HRF leads to loss of power that cannot be accounted for by adding time and dispersion derivatives. We provide a first estimate of the cortical dog HRF resulting in significant activation increase in comparison to the human HRF and validated our results using an independent task. We hope that our findings spark new research that might challenge or add to our results. To increase transparency, we applied open-science practices throughout, and hope this will motivate and facilitate future investigations by other labs, leading to a joint effort to improve detection power in canine neuroimaging research.

## 5 Conflicts of interest

The authors declare no competing financial interests.

## 6 Acknowledgements

We thank Lukas Lengersdorff for his helpful comments on the analysis plan, Morris Krainz for his help collecting the data and all the dogs and their caregivers for taking part in our study. This project was supported by the Austrian Science Fund (FWF): W1262-B29 and by the Vienna Science and Technology Fund (WWTF), the City of Vienna and ithuba Capital AG through project CS18-012. The funders had no role in study design, data collection and analysis, decision to publish, or preparation of the manuscript.

## References

Aguirre, G. K., Komáromy, A. M., Cideciyan, A. V, Brainard, D. H., Aleman, T. S., Roman, A. J., … Jacobson, S. G. (2007). Canine and Human Visual Cortex Intact and Responsive Despite Early Retinal Blindness from RPE65 Mutation. PLoS Medicine, 4(6), e230. https://doi.org/10.1371/journal.pmed.0040230

Aguirre, G. K., Zarahn, E., & D’Esposito, M. (1998). The Variability of Human, BOLD Hemodynamic Responses. NeuroImage, 8(4), 360–369. https://doi.org/10.1006/nimg.1998.0369

Andics, A., Gábor, A., Gácsi, M., Faragó, T., Szabó, D., & Miklósi, Á. (2016). Neural mechanisms for lexical processing in dogs. Science, 353(6303), 1030–1032. https://doi.org/10.1126/science.aaf3777

Andics, A., Gácsi, M., Faragó, T., Kis, A., & Miklósi, Á. (2014). Voice-Sensitive Regions in the Dog and Human Brain Are Revealed by Comparative fMRI. Current Biology, 24(5), 574–578. https://doi.org/10.1016/J.CUB.2014.01.058

Andics, A., & Miklósi, Á. (2018). Neural processes of vocal social perception: Dog-human comparative fMRI studies. Neuroscience & Biobehavioral Reviews, 85, 54–64. https://doi.org/10.1016/j.neubiorev.2017.11.017

Arichi, T., Fagiolo, G., Varela, M., Melendez-Calderon, A., Allievi, A., Merchant, N., … Edwards, A. D. (2012). Development of BOLD signal hemodynamic responses in the human brain. NeuroImage, 63(2), 663–673. https://doi.org/10.1016/j.neuroimage.2012.06.054

Aulet, L. S., Chiu, V. C., Prichard, A., Spivak, M., Lourenco, S. F., & Berns, G. S. (2019). Canine sense of quantity: Evidence for numerical ratio-dependent activation in parietotemporal cortex. Biology Letters, 15(12). https://doi.org/10.1098/rsbl.2019.0666

Baumann, S., Griffiths, T. D., Rees, A., Hunter, D., Sun, L., & Thiele, A. (2010). Characterisation of the BOLD response time course at different levels of the auditory pathway in non-human primates. NeuroImage, 50(3), 1099–1108. https://doi.org/10.1016/j.neuroimage.2009.12.103

Berns, G. S., Brooks, A. M., & Spivak, M. (2012). Functional MRI in Awake Unrestrained Dogs. PLoS ONE, 7(5), e38027. https://doi.org/10.1371/journal.pone.0038027

Berns, G. S., Brooks, A. M., & Spivak, M. (2015). Scent of the familiar: An fMRI study of canine brain responses to familiar and unfamiliar human and dog odors. Behavioural Processes, 110, 37–46. https://doi.org/10.1016/J.BEPROC.2014.02.011

Berns, G. S., Brooks, A. M., Spivak, M., & Levy, K. (2017). Functional MRI in Awake Dogs Predicts Suitability for Assistance Work. Scientific Reports, 7, 43704. https://doi.org/10.1038/srep43704

Berns, G. S., Brooks, A., & Spivak, M. (2013). Replicability and Heterogeneity of Awake Unrestrained Canine fMRI Responses. PLoS ONE, 8(12), e81698. https://doi.org/10.1371/journal.pone.0081698

Berns, G. S., & Cook, P. F. (2016). Why Did the Dog Walk Into the MRI? Current Directions in Psychological Science, 25(5), 363–369. https://doi.org/10.1177/0963721416665006

Boynton, G. M., Engel, S. A., Glover, G. H., & Heeger, D. J. (1996). Linear systems Analysis of Functional Magnetic Resonance Imaging in Human V1. Journal of Neuroscience, 16(13), 4207–4221. https://doi.org/10.1523/jneurosci.16-13-04207.1996

Bunford, N., Andics, A., Kis, A., Miklósi, Á., & Gácsi, M. (2017). Canis familiaris As a Model for Non-Invasive Comparative Neuroscience. Trends in Neurosciences, 40(7), 438–452. https://doi.org/10.1016/J.TINS.2017.05.003

Button, K. S., Ioannidis, J. P. A., Mokrysz, C., Nosek, B. A., Flint, J., Robinson, E. S. J., & Munafò, M. R. (2013). Power failure: why small sample size undermines the reliability of neuroscience. Nature Reviews Neuroscience, 14(5), 365–376. https://doi.org/10.1038/nrn3475

Calhoun, V. D., Stevens, M. C., Pearlson, G. D., & Kiehl, K. A. (2004). fMRI analysis with the general linear model: removal of latency-induced amplitude bias by incorporation of hemodynamic derivative terms. NeuroImage, 22(1), 252–257. https://doi.org/10.1016/j.neuroimage.2003.12.029

Carp, J. (2012a). On the plurality of (methodological) worlds: estimating the analytic flexibility of fMRI experiments. Frontiers in Neuroscience, 6, 149. https://doi.org/10.3389/fnins.2012.00149

Carp, J. (2012b). The secret lives of experiments: Methods reporting in the fMRI literature. NeuroImage, 63(1), 289–300. https://doi.org/10.1016/j.neuroimage.2012.07.004

Chen, X., Tong, C., Han, Z., Zhang, K., Bo, B., Feng, Y., & Liang, Z. (2020). Sensory evoked fMRI paradigms in awake mice. NeuroImage, 204. https://doi.org/10.1016/j.neuroimage.2019.116242

Cook, P. F., Prichard, A., Spivak, M., & Berns, G. S. (2016). Awake canine fMRI predicts dogs’ preference for praise vs food. Social Cognitive and Affective Neuroscience, 11(12), 1853–1862. https://doi.org/10.1093/scan/nsw102

Cook, P. F., Prichard, A., Spivak, M., & Berns, G. S. (2018). Jealousy in dogs? Evidence from brain imaging. Animal Sentience, 22(1), 1–14.

Cook, P. F., Spivak, M., & Berns, G. S. (2014). One pair of hands is not like another: caudate BOLD response in dogs depends on signal source and canine temperament. PeerJ, 2:e596; https://doi.org/10.7717/peerj.596

Cremers, H. R., Wager, T. D., & Yarkoni, T. (2017). The relation between statistical power and inference in fMRI. PLOS ONE, 12(11), e0184923. https://doi.org/10.1371/journal.pone.0184923

Cuaya, L. V., Hernández-Pérez, R., & Concha, L. (2016). Our Faces in the Dog’s Brain: Functional Imaging Reveals Temporal Cortex Activation during Perception of Human Faces. PLOS ONE, 11(3), e0149431. https://doi.org/10.1371/journal.pone.0149431

Czeibert, K., Andics, A., Petneházy, Ö., & Kubinyi, E. (2019). A detailed canine brain label map for neuroimaging analysis. Biologia Futura, 70(2), 112–120. https://doi.org/10.1556/019.70.2019.14

Datta, R., Lee, J., Duda, J., Avants, B. B., Vite, C. H., Tseng, B., … Aguirre, G. K. (2012). A Digital Atlas of the Dog Brain. PLoS ONE, 7(12), e52140. https://doi.org/10.1371/journal.pone.0052140

De Zwart, J. A., Silva, A. C., Van Gelderen, P., Kellman, P., Fukunaga, M., Chu, R., … Duyn, J. H. (2005). Temporal dynamics of the BOLD fMRI impulse response. NeuroImage, 24(3), 667–677. https://doi.org/10.1016/j.neuroimage.2004.09.013

DeFelipe, J. (2011). The evolution of the brain, the human nature of cortical circuits, and intellectual creativity. Frontiers in Neuroanatomy, 5, 1–29. https://doi.org/10.3389/fnana.2011.00029

Dilks, D. D., Cook, P. F., Weiller, S. K., Berns, H. P., Spivak, M., & Berns, G. S. (2015). Awake fMRI reveals a specialized region in dog temporal cortex for face processing. PeerJ, 3:e1115. https://doi.org/10.7717/peerj.1115

Fitch, W. T., Huber, L., & Bugnyar, T. (2010). Social Cognition and the Evolution of Language: Constructing Cognitive Phylogenies. Neuron, 65(6), 795–814. https://doi.org/10.1016/J.NEURON.2010.03.011

Ford, J. M., Johnson, M. B., Whitfield, S. L., Faustman, W. O., & Mathalon, D. H. (2005). Delayed hemodynamic responses in schizophrenia. NeuroImage, 26(3), 922–931. https://doi.org/10.1016/j.neuroimage.2005.03.001

Friston, K. J., Fletcher, P., Josephs, O., Holmes, A., Rugg, M. D., & Turner, R. (1998). Event-related fMRI: Characterizing Differential Responses. NeuroImage, 7, 30–40. https://doi.org/10.1006/nimg.1997.0306

Friston, K. J., Holmes, A. P., Poline, J.-B., Grasby, P. J., Williams, S. C. R., Frackowiak, R. S. J., & Turner, R. (1995). Analysis of fMRI Time-Series Revisited. NeuroImage, 2, 45–53. https://doi.org/10.1006/nimg.1995.1007

Friston, K. J., Jezzard, P., & Turner, R. (1994). Analysis of functional MRI timeseries. Human Brain Mapping, 1(2), 153–171. https://doi.org/10.1002/hbm.460010207

Friston, K. J., Josephs, O., Rees, G., & Turner, R. (1998). Nonlinear event-related responses in fMRI. Magnetic Resonance in Medicine, 39(1), 41–52. https://doi.org/10.1002/mrm.1910390109

Glover, G. H. (1999). Deconvolution of impulse response in event-related BOLD fMRI. NeuroImage, 9(4), 416–429. https://doi.org/10.1006/nimg.1998.0419

Goense, J. B. M., & Logothetis, N. K. (2008). Neurophysiology of the BOLD fMRI Signal in Awake Monkeys. Current Biology, 18(9), 631–640. https://doi.org/10.1016/j.cub.2008.03.054

Handwerker, D. A., Ollinger, J. M., & D’Esposito, M. (2004). Variation of BOLD hemodynamic responses across subjects and brain regions and their effects on statistical analyses. NeuroImage, 21(4), 1639–1651. https://doi.org/10.1016/j.neuroimage.2003.11.029

Hecht, E. E., Smaers, J. B., Dunn, W. J., Kent, M., Preuss, T. M., & Gutman, D. A. (2019). Significant Neuroanatomical Variation Among Domestic Dog Breeds. Journal of Neuroscience, 39(39), 0303–0319. https://doi.org/10.1523/JNEUROSCI.0303-19.2019

Henson, R. N. A., Price, C. J., Rugg, M. D., Turner, R., & Friston, K. J. (2002). Detecting Latency Differences in Event-Related BOLD responses: Application to Words versus Nonwords and Initial versus Repeated Face Presentations. NeuroImage, 15(1), 83–97. https://doi.org/10.1006/nimg.2001.0940

Hernández-Pérez, R., Concha, L., & Cuaya, L. V. (2018). Decoding Human Emotional Faces in the Dog’s Brain. BioRxiv. https://doi.org/10.1101/134080

Horschler, D. J., Hare, B., Call, J., Kaminski, J., Miklósi, Á., & MacLean, E. L. (2019). Absolute brain size predicts dog breed differences in executive function. Animal Cognition, 22(2), 187–198. https://doi.org/10.1007/s10071-018-01234-1

Huber, L., & Lamm, C. (2017). Understanding dog cognition by functional magnetic resonance imaging. Learning & Behavior, 45(2), 101–102. https://doi.org/10.3758/s13420-017-0261-6

Ioannidis, J. P. A. (2005). Why Most Published Research Findings Are False. PLoS Medicine, 2(8), e124. https://doi.org/10.1371/journal.pmed.0020124

Jia, H., Pustovyy, O. M., Waggoner, P., Beyers, R. J., Schumacher, J., Wildey, C., … Deshpande, G. (2014). Functional MRI of the Olfactory System in Conscious Dogs. PLoS ONE, 9(1), e86362. https://doi.org/10.1371/journal.pone.0086362

Jia, H., Pustovyy, O. M., Wang, Y., Waggoner, P., Beyers, R. J., Schumacher, J., … Deshpande, G. (2016). Enhancement of Odor-Induced Activity in the Canine Brain by Zinc Nanoparticles: A Functional MRI Study in Fully Unrestrained Conscious Dogs. Chemical Senses, 41, 53–67. https://doi.org/10.1093/chemse/bjv054

Karl, S., Boch, M., Virányi, Z., Lamm, C., & Huber, L. (2019). Training pet dogs for eye-tracking and awake fMRI. Behavior Research Methods, 1–19. https://doi.org/10.3758/s13428-019-01281-7

Kilkenny, C., Browne, W. J., Cuthill, I. C., Emerson, M., & Altman, D. G. (2010). Improving Bioscience Research Reporting: The ARRIVE Guidelines for Reporting Animal Research. PLoS Biology, 8(6), e1000412. https://doi.org/10.1371/journal.pbio.1000412

Koyama, M., Hasegawa, I., Osada, T., Adachi, Y., Nakahara, K., & Miyashita, Y. (2004). Functional Magnetic Resonance Imaging Of Macaque Monkeys Performing Visually Guided Saccade Tasks: Comparison of Cortical Eye Fields with Humans. Neuron, 41(5), 795–807. https://doi.org/10.1016/S0896-6273(04)00047-9

Lambers, H., Segeroth, M., Albers, F., Wachsmuth, L., van Alst, T. M., & Faber, C. (2020). A cortical rat hemodynamic response function for improved detection of BOLD activation under common experimental conditions. NeuroImage, 208. https://doi.org/10.1016/j.neuroimage.2019.116446

Langley, J. N., & Grünbaum, A. S. (1890). On the degeneration resulting from removal of the cerebral cortex and corpora striata in the dog. The Journal of Physiology, 11, 606.

Lewis, L. D., Setsompop, K., Rosen, B. R., & Polimeni, J. R. (2018). Stimulus-dependent hemodynamic response timing across the human subcortical-cortical visual pathway identified through high spatiotemporal resolution 7T fMRI. NeuroImage, 181, 279–291. https://doi.org/10.1016/j.neuroimage.2018.06.056

Lindquist, M. A., Meng Loh, J., Atlas, L. Y., & Wager, T. D. (2009). Modeling the hemodynamic response function in fMRI: Efficiency, bias and mis-modeling. NeuroImage, 45(1). https://doi.org/10.1016/j.neuroimage.2008.10.065

Liu, X., Tian, R., Zuo, Z., Zhao, H., Wu, L., Zhuo, Y., … Chen, L. (2020). A highresolution MRI brain template for adult Beagle. Magnetic Resonance Imaging, 68, 148–157. https://doi.org/10.1016/j.mri.2020.01.003

Logothetis, N. K., Pauls, J., Augath, M., Trinath, T., & Oeltermann, A. (2001). Neurophysiological investigation of the basis of the fMRI signal. Nature, 412(6843), 150–157. https://doi.org/10.1038/35084005

Marquis, D. G. (1934). Effects of removal of the visual cortex in mammals, with observations on the retention of light discrimination in dogs. Association for Research in Nervous and Mental Disease, 558–592.

Nakahara, K., Hayashi, T., Konishi, S., & Miyashita, Y. (2002). Functional MRI of Macaque Monkeys Performing a Cognitive Set-Shifting Task. Science, 295(5559), 1532–1536. https://doi.org/10.1126/science.1067653

Nichols, T. E., Das, S., Eickhoff, S. B., Evans, A. C., Glatard, T., Hanke, M., … Yeo, B. T. T. (2017). Best practices in data analysis and sharing in neuroimaging using MRI. Nature Neuroscience, 20(3), 299–303. https://doi.org/10.1038/nn.4500

Nitzsche, B., Boltze, J., Ludewig, E., Flegel, T., Schmidt, M. J., Seeger, J., … Schulze, S. (2019). A stereotaxic breed-averaged, symmetric T2w canine brain atlas including detailed morphological and volumetrical data sets. NeuroImage, 187, 93–103. https://doi.org/10.1016/j.neuroimage.2018.01.066

Orban, G. A., Van Essen, D., & Vanduffel, W. (2004, July). Comparative mapping of higher visual areas in monkeys and humans. Trends in Cognitive Sciences, Vol. 8, pp. 315–324. https://doi.org/10.1016/j.tics.2004.05.009

Patel, G. H., Cohen, A. L., Baker, J. T., Snyder, L. H., & Corbetta, M. (2018). Comparison of stimulus-evoked BOLD responses in human and monkey visual cortex. BioRxiv, 345330. https://doi.org/10.1101/345330

Poldrack, R. A., Baker, C. I., Durnez, J., Gorgolewski, K. J., Matthews, P. M., Munafò, M. R., … Yarkoni, T. (2017). Scanning the horizon: towards transparent and reproducible neuroimaging research. Nature Reviews Neuroscience, 18(2), 115–126. https://doi.org/10.1038/nrn.2016.167

Poldrack, R. A., Fletcher, P. C., Henson, R. N., Worsley, K. J., Brett, M., & Nichols, T. E. (2008). Guidelines for reporting an fMRI study. NeuroImage, 40(2), 409–414. https://doi.org/10.1016/J.NEUROIMAGE.2007.11.048

Power, J. D., Barnes, K. A., Snyder, A. Z., Schlaggar, B. L., & Petersen, S. E. (2012). Spurious but systematic correlations in functional connectivity MRI networks arise from subject motion. NeuroImage, 59(3), 2142–2154. https://doi.org/10.1016/J.NEUROIMAGE.2011.10.018

Power, J. D., Mitra, A., Laumann, T. O., Snyder, A. Z., Schlaggar, B. L., & Petersen, S. E. (2014). Methods to detect, characterize, and remove motion artifact in resting state fMRI. NeuroImage, 84, 320–341. https://doi.org/10.1016/j.neuroimage.2013.08.048

Prichard, A., Chhibber, R., Athanassiades, K., Spivak, M., & Berns, G. S. (2018). Fast neural learning in dogs: A multimodal sensory fMRI study. Scientific Reports, 8, 14614. https://doi.org/10.1038/s41598-018-32990-2

Prichard, A., Chhibber, R., King, J., Athanassiades, K., Spivak, M., & Berns, G. S. (2019). Decoding Odor Mixtures in the Dog Brain: An Awake fMRI Study. BioRxiv, 754374. https://doi.org/10.1101/754374

Prichard, A., Cook, P. F., Spivak, M., Chhibber, R., & Berns, G. S. (2018). Awake fMRI Reveals Brain Regions for Novel Word Detection in Dogs. Frontiers in Neuroscience, 12, 737. https://doi.org/10.3389/fnins.2018.00737

Roth, G., & Dicke, U. (2005). Evolution of the brain and intelligence. Trends in Cognitive Sciences, 9(5), 250–257. https://doi.org/10.1016/j.tics.2005.03.005

Schoenebeck, J. J., & Ostrander, E. A. (2013). The Genetics of Canine Skull Shape Variation. Genetics, 193(2), 317–325. https://doi.org/10.1534/genetics.112.145284

Silva, A. C., Koretsky, A. P., & Duyn, J. H. (2007). Functional MRI impulse response for BOLD and CBV contrast in rat somatosensory cortex. Magnetic Resonance in Medicine, 57(6), 1110–1118. https://doi.org/10.1002/mrm.21246

Simmons, J. P., Nelson, L. D., & Simonsohn, U. (2011). False-Positive Psychology: Undisclosed Flexibility in Data Collection and Analysis Allows Presenting Anything as Significant. Psychological Science, 22(11), 1359–1366. https://doi.org/10.1177/0956797611417632

Sladky, R., Friston, K. J., Tröstl, J., Cunnington, R., Moser, E., & Windischberger, C. (2011). Slice-timing effects and their correction in functional MRI. NeuroImage, 58(2), 588–594. https://doi.org/10.1016/J.NEUROIMAGE.2011.06.078

Strassberg, L. R., Waggoner, L. P., Deshpande, G., & Katz, J. S. (2019). Training Dogs for Awake, Unrestrained Functional Magnetic Resonance Imaging. Journal of Visualized Experiments: JoVE, (152). https://doi.org/10.3791/60192

Szabó, D., Gábor, A., Gácsi, M., Faragó, T., Kubinyi, E., Miklósi, Á., & Andics, A. (2020). On the Face of It: No Differential Sensitivity to Internal Facial Features in the Dog Brain. Frontiers in Behavioral Neuroscience, 14, 25. https://doi.org/10.3389/fnbeh.2020.00025

Thompkins, A. M., Deshpande, G., Waggoner, P., & Katz, J. S. (2016). Functional Magnetic Resonance Imaging of the Domestic Dog: Research, Methodology, and Conceptual Issues. Comparative Cognition & Behavior Reviews, 11, 63–82. https://doi.org/10.3819/ccbr.2016.110004

Thompkins, A. M., Ramaiahgari, B., Zhao, S., Gotoor, S. S. R., Waggoner, P., Denney, T. S., … Katz, J. S. (2018). Separate brain areas for processing human and dog faces as revealed by awake fMRI in dogs (Canis familiaris). Learning and Behavior, 46(4), 561–573. https://doi.org/10.3758/s13420-018-0352-z

Tootell, R. B. H., Tsao, D., & Vanduffel, W. (2003, May 15). Neuroimaging Weighs In: Humans Meet Macaques in “Primate” Visual Cortex. Journal of Neuroscience, Vol. 23, pp. 3981–3989. https://doi.org/10.1523/jneurosci.23-10-03981.2003

Uemura, E. E. (2015). Fundamentals of Canine Neuroanatomy and Neurophysiology. Ames, Iowa: John Wiley & Sons, Inc.

Upham, N. S., Esselstyn, J. A., & Jetz, W. (2019). Inferring the mammal tree: Species-level sets of phylogenies for questions in ecology, evolution, and conservation. PLoS Biology, 17(12), e3000494. https://doi.org/10.1371/journal.pbio.3000494

Willis, C. K. R., Quinn, R. P., McDonell, W. M., Gati, J., Parent, J., & Nicolle, D. (2001). Functional MRI as a tool to assess vision in dogs: The optimal anesthetic. Veterinary Ophthalmology, 4(4), 243–253. https://doi.org/10.1046/j.1463-5216.2001.00183.x

Willis, C. K. R., Quinn, R. P., McDonell, W. M., Gati, J., Partlow, G., & Vilis, T. (2001). Functional MRI activity in the thalamus and occipital cortex of anesthetized dogs induced by monocular and binocular stimulation. Canadian Journal of Veterinary Research = Revue Canadienne de Recherche Veterinaire, 65(3), 188–195. Retrieved from http://www.ncbi.nlm.nih.gov/pubmed/11480525

Wing, K. G., & Smith, K. U. (1942). The role of the optic cortex in the dog in the determination of the functional properties of conditioned reactions to light. Journal of Experimental Psychology, 31(6), 478–496.

Worsley, K. J., & Friston, K. J. (1995). Analysis of fMRI Time-Series Revisited — Again. NeuroImage, 2, 173–181. https://doi.org/10.1006/nimg.1995.1023

Yáñez, I. B., Muñoz, A., Contreras, J., Gonzalez, J., Rodriguez-Veiga, E., & DeFelipe, J. (2005). Double bouquet cell in the human cerebral cortex and a comparison with other mammals. Journal of Comparative Neurology, 486(4), 344–360. https://doi.org/10.1002/cne.20533

Yushkevich, P. A., Piven, J., Hazlett, H. C., Smith, R. G., Ho, S., Gee, J. C., & Gerig, G. (2006). User-guided 3D active contour segmentation of anatomical structures: Significantly improved efficiency and reliability. NeuroImage, 31(3), 1116–1128. https://doi.org/10.1016/J.NEUROIMAGE.2006.01.015

